# Rho and F-actin self-organize within an artificial cell cortex

**DOI:** 10.1101/2021.04.09.438460

**Authors:** Jennifer Landino, Marcin Leda, Ani Michaud, Zachary T. Swider, Mariah Prom, Christine M. Field, William M. Bement, Anthony G. Vecchiarelli, Andrew B. Goryachev, Ann L. Miller

## Abstract

The cell cortex, comprised of the plasma membrane and underlying cytoskeleton, undergoes dynamic reorganizations during a variety of essential biological processes including cell adhesion, cell migration, and cell division^1,2^. During cell division and cell locomotion, for example, waves of filamentous-actin (F-actin) assembly and disassembly develop in the cell cortex in a process termed “cortical excitability”^3–7^. In developing frog and starfish embryos, cortical excitability is generated through coupled positive and negative feedback, with rapid activation of Rho-mediated F-actin assembly followed in space and time by F-actin-dependent inhibition of Rho^8,9^. These feedback loops are proposed to serve as a mechanism for amplification of active Rho signaling at the cell equator to support furrowing during cytokinesis, while also maintaining flexibility for rapid error correction in response to movement of the mitotic spindle during chromosome segregation^10^. In this paper, we develop an artificial cortex based on *Xenopus* egg extract and supported lipid bilayers (SLBs), to investigate cortical Rho and F-actin dynamics^11^. This reconstituted system spontaneously develops two distinct dynamic patterns: singular excitable Rho and F-actin waves and non-traveling oscillatory Rho and F-actin patches. Both types of dynamic patterns have properties and dependencies similar to the cortical excitability previously characterized *in vivo*^9^. These findings directly support the longstanding speculation that the cell cortex is a self-organizing structure and present a novel approach for investigating mechanisms of Rho-GTPase-mediated cortical dynamics.

**Highlights:**

- An artificial cell cortex comprising *Xenopus* egg extract on a supported lipid bilayer self-organizes into complex, dynamic patterns of active Rho and F-actin
- We identified two types of reconstituted cortical dynamics – excitable waves and coherent oscillations
- Reconstituted waves and oscillations require Rho activity and F-actin polymerization

## Results and Discussion

### Reconstituted cortex generates excitable waves of active Rho and F-actin

A cell-free system comprised of *Xenopus* egg extract containing intact F-actin and artificial centrosomes has been previously shown to reconstitute cortical cytokinetic patterning. Microtubule-dependent signaling localizes active Rho within an artificial cell cortex formed atop SLBs^11^. Here we endeavor to reconstitute self-organized cortical Rho patterning, independent of exogenous signals. Specifically, we adapted a similar cell-free system to investigate active Rho and F-actin waves that have previously only been observed in live cells^9^. To visualize cortical Rho and actin dynamics, we added *Xenopus* egg extract containing recombinant probes for active Rho (Rho binding domain of Rhotekin (rGBD)) and F-Actin (Utrophin calponin homology domain (UtrCH)) to SLBs designed to mimic the inner leaflet of the animal cell membrane^11–13^. We imaged active Rho and F-actin associated with SLBs using TIRF microscopy (Supplemental Figure 1A). Using this approach, we identified two distinct types of self-organized cortical dynamics: 1) an excitable, propagating wave front of active Rho and F-actin polymerization and 2) coherent, oscillatory dynamics of active Rho and F-actin with the SLBs.

Immediately after adding *Xenopus* extract to the SLBs, we observed the rapid selforganization of active Rho and F-actin patterning in the form of a dramatic propagation of a solitary excitable wave (Figure 1A and 1B). This remarkable phenomenon manifested as a sudden large-amplitude increase in Rho activity in the form of several discrete maxima (Figure 1A and 1C, white arrows) that rapidly spread and transformed into irregular circular waves that annihilated upon collision with each other (Figure 1A and 1C, yellow arrowheads, Supplemental Movie 1). Two-color imaging with high temporal resolution demonstrated that the Rho activity wave is closely followed by the actin polymerization wave (Figure 1B and 1D). This excitable behavior started and ended within 140 ± 38 seconds (mean ± standard deviation, n = 5 experiments) after adding *Xenopus* egg extract to the SLBs, occurring just once per reconstituted cortex image acquisition sequence (Supplemental Movie 2). Thus, the self-organized excitable wave observed in the reconstituted system was solitary, and the nature of its initiation remains unclear.

**Figure 1:**
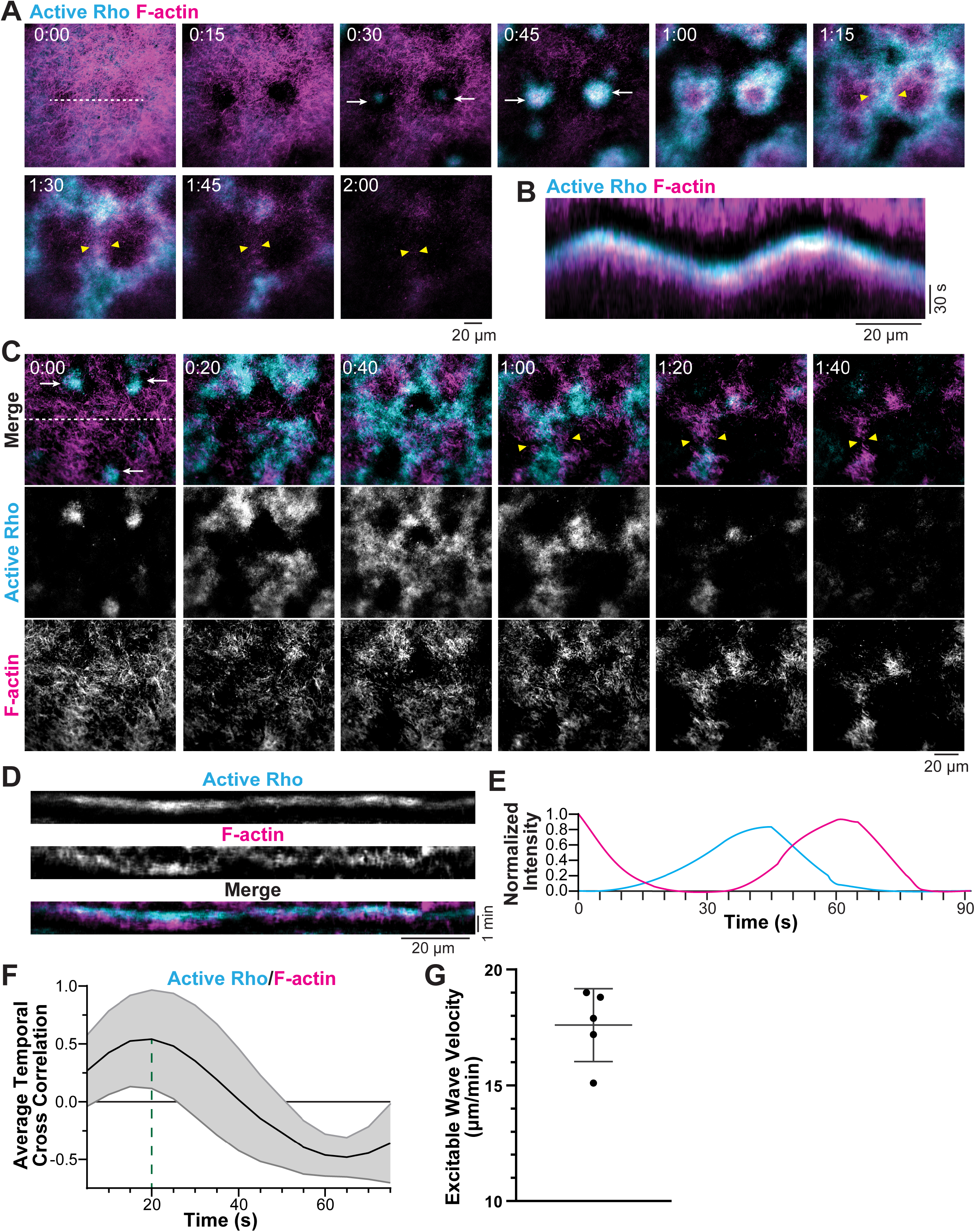
Artificial cortex supports the self-organization of excitable waves of active Rho and F-actin. A) Micrographs (30s difference subtraction) of traveling excitable waves of active Rho (cyan) and F-actin (magenta) that originate from discrete maxima (white arrows). Yellow arrowheads indicate wave fronts that annihilate on collision. Time is in minutes:seconds. Dashed line represents region used to generate kymograph. B) Kymographs of excitable waves of active Rho (cyan) and F-actin (magenta) generated from a 10 pixel wide line shown in (A). C) Micrographs of excitable waves of active Rho (cyan) and F-actin (magenta) used for analysis (E-G). White arrows represent wave origins, and yellow arrowheads indicate wave annihilations. Time is in minutes:seconds. Dashed line represents region used to generate kymograph. D) Kymographs of excitable waves of active Rho (cyan) and F-actin (magenta) generated from a 10 pixel wide line shown in (C). E) Normalized time series of intensities of active Rho and F-actin spatially averaged over a representative 20×20 pixel box selected from the field of view (see Supplemental Figure 1B). Moving time average of three frames. F) Temporal cross correlation between active Rho and F-actin intensities. Dashed line indicates the time shift of 20.7 s. G) Excitable wave velocity, 17.6 ± 1.6 microns/minute (mean ± standard deviation), was measured over 50-60 s for 18 wave fronts across 5 experiments. Each dot represents the average velocity for one experiment.

To specifically visualize and quantify wave dynamics, we used temporal difference subtraction to computationally remove static active Rho and F-actin signal (Figure 1C, Supplemental Figure 1B)^9^. We then divided the field of view into a grid of 20×20 pixel boxes and plotted the box-averaged and normalized fluorescence intensity of active Rho and F-actin in an individual box over time (Figure 1E). This enabled us to measure the cross-correlation between the dynamics of active Rho and F-actin (Figure 1F, Supplemental Figure 2B). Across multiple experiments, the time shift between the maxima of Rho activity and F-actin polymerization within excitable waves was 22.6 ± 6.3 seconds (mean ± standard deviation, n = 5 experiments, 3027 boxes, Supplemental Figure 2C). We also quantified the velocity of excitable waves, using kymographs to track the position of the wave front over time. Excitable active Rho waves traveled at 17.6 ± 1.6 μm/minute (mean ± standard deviation, n = 5 experiments, 151 wave fronts, Figure 1G, Supplemental Figure 2D), which is similar to the velocity of cortical waves in *Xenopus* blastomeres (13.5 μm/minute, Supplemental Figure 3E)^9^.

What could explain the remarkable solitary occurrence of excitable waves in our experiments? We considered the possibility that in the cell-free system, there may be limiting components, which are consumed during the first excitable wave, preventing the continued propagation of additional excitable waves. We tested this possibility by increasing the volume of the extract relative to the surface area of the SLBs, supplementing GTP, or increasing the relative proportion of PI(4,5)P_2_ in the lipid content of the SLBs. Surprisingly, none of these adjustments changed the solitary nature of excitable waves (data not shown).

### Active Rho and F-actin coherently oscillate in a cell-free reconstituted system

In addition to excitable waves, the reconstituted cortex self-organized into oscillatory, coherent pulses of active Rho and F-actin which generally occurred after the appearance of an excitable wave, appearing 13 ± 9 minutes (mean ± standard deviation, n = 8 experiments) after the extract is added to the SLB. These oscillations exhibit a defined temporal period, but do not result in spatially propagating waves instead forming discrete patches on the SLB (Figure 2A, 2B, and 2C, Supplemental Movie 3). In addition to the oscillatory active Rho and F-actin, we also observed prominent static fluorescence signals for active Rho and F-actin that progressively accumulate on the bilayer, partially masking the oscillatory dynamics of active Rho and F-actin on the SLB (Supplemental Figure 3A). Again, we used temporal difference subtraction to computationally remove static signals prior to quantifying oscillatory dynamics (Supplemental Figure 1B)^9^.

**Figure 2:**
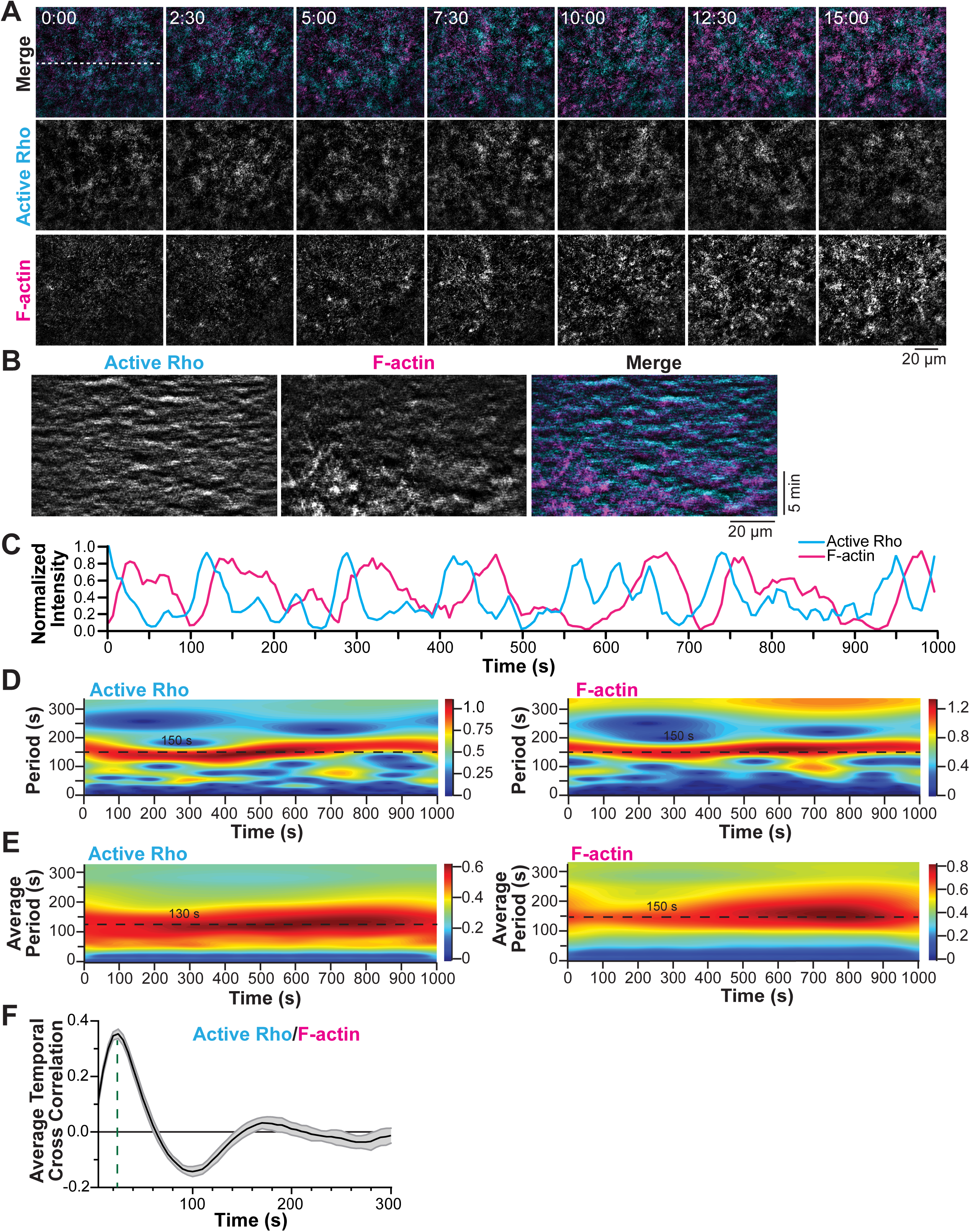
Active Rho and F-actin coherently oscillate within an artificial cortex. A) Reconstituted oscillatory dynamics of active Rho (cyan) and F-actin (magenta). Time is indicated in minutes:seconds. Dashed line represents the region used to generate kymographs. B) Kymographs of oscillatory dynamics of active Rho (cyan) and F-actin (magenta) generated from a 10 pixel wide line shown in panel (A). C) Normalized time series of intensities of active Rho and F-actin spatially averaged over a representative 20×20 pixel box selected from the field of view. Moving time average of three frames. D) Morlet power spectra showing the time-dependent periodicity of the active Rho and F-actin dynamics computed for the same 20×20 pixel box as in (C). Dashed lines indicate dominant oscillatory periods. E) Morlet power spectra of the active Rho and F-actin dynamics averaged over all boxes constituting the whole field of view shown in (A). Dashed lines indicate dominant oscillatory periods. F) Temporal cross correlation between active Rho and F-actin fluorescence signals. Dashed line indicates the time shift of 25 s between the two oscillations.

The temporal dynamics of active Rho and F-actin were analyzed using a Morlet wavelet transform that was applied to respective fluorescence signals spatially averaged over individual boxes (Figure 2D)^9,14^. This local analysis revealed complex, spatially and temporally heterogeneous behavior with short oscillatory stretches of varying period appearing and vanishing throughout the experiment. Remarkably, spatial averaging of local Morlet spectra revealed a robust dominant oscillatory pattern with a reproducible period indicating that the accumulation of static Rho and F-actin on the SLB does not significantly affect cortical oscillations (Figure 2E). Across multiple experiments, we determined the period of active Rho oscillations to be 131 ± 8.6 seconds (mean ± standard deviation, n=8 experiments, 2453 boxes, Supplemental Figure 3C). This is comparable to the period of waves of active Rho previously characterized in developing *Xenopus* embryos (80-120 seconds)^9^. Using temporal cross-correlation analysis (Figure 2F), we determined that oscillatory F-actin peaked 22.5 ± 4.1 seconds (mean ± standard deviation, n=8 experiments, 5494 boxes) after the respective maxima of active Rho (Supplemental Figure 3D), which is well-matched to the time shift for excitable waves of active Rho and F-actin (Supplemental Figure 3E).

We also investigated whether these coherent oscillations could develop into propagating excitable waves, similar to the self-organized cortical waves observed immediately after adding extract to the SLB. To explore this possibility, we sought to extend the phase of the extract cell cycle in which the cortex can support excitability (low Cdk1 activity, I-phase)^15^ and prevent the extract from transitioning into M-phase where actomyosin is contractile^11,12,15,16^. Blocking cell cycle progression with pharmacological inhibitors (cycloheximide, a protein synthesis inhibitor) or RO-3306 (a Cdk1 inhibitor)) did not promote the transition from oscillatory to excitable dynamics. Rather, we sometimes observed that coherent oscillations of active Rho and F-actin increased in size over time, but the fundamental characteristics of their dynamics did not change (data not shown).

Our analysis of both excitable and oscillatory Rho and F-actin patterns demonstrates that the artificial cortex successfully reconstitutes cortical dynamics with characteristics similar to those described *in vivo* (Supplemental Figure 3E)^9^. Moreover, the emergence of cortical patterns, in the form of Rho and F-actin waves and oscillations, in this cell-free system reveals that the artificial cortex can self-organize independently of exogenous signaling from sources such as the nucleus or centrosomes which are not present in our experiments.

### Reconstituted Rho and F-actin oscillations and waves require Rho activity

Given the quantifiable similarities between the reconstituted Rho oscillations and *in vivo* cortical waves, we next asked whether the molecular mechanism that underlies cortical Rho dynamics was conserved in the reconstituted system. We first investigated whether Rho activity is required for the dynamic Rho and F-actin patterning observed on SLBs. To test this, we treated the extract with a Rho activity inhibitor, C3 transferase, prior to adding the extract to the SLB^17^. Using two concentrations of C3 transferase (33 μg/ml and 100 μg/ml), we found that treatment with C3 prevented both oscillatory and excitable dynamics of Rho and F-actin (Figure 3A, 3B, and 3C, Supplemental Movie 4)^18^. These data indicate that Rho activity is essential for the observed dynamic patterning in the reconstituted system. This result is in agreement with the work done in *Xenopus* and starfish embryos^9^, and suggests that Rho-mediated positive feedback initiates cortical excitability and oscillations.

**Figure 3:**
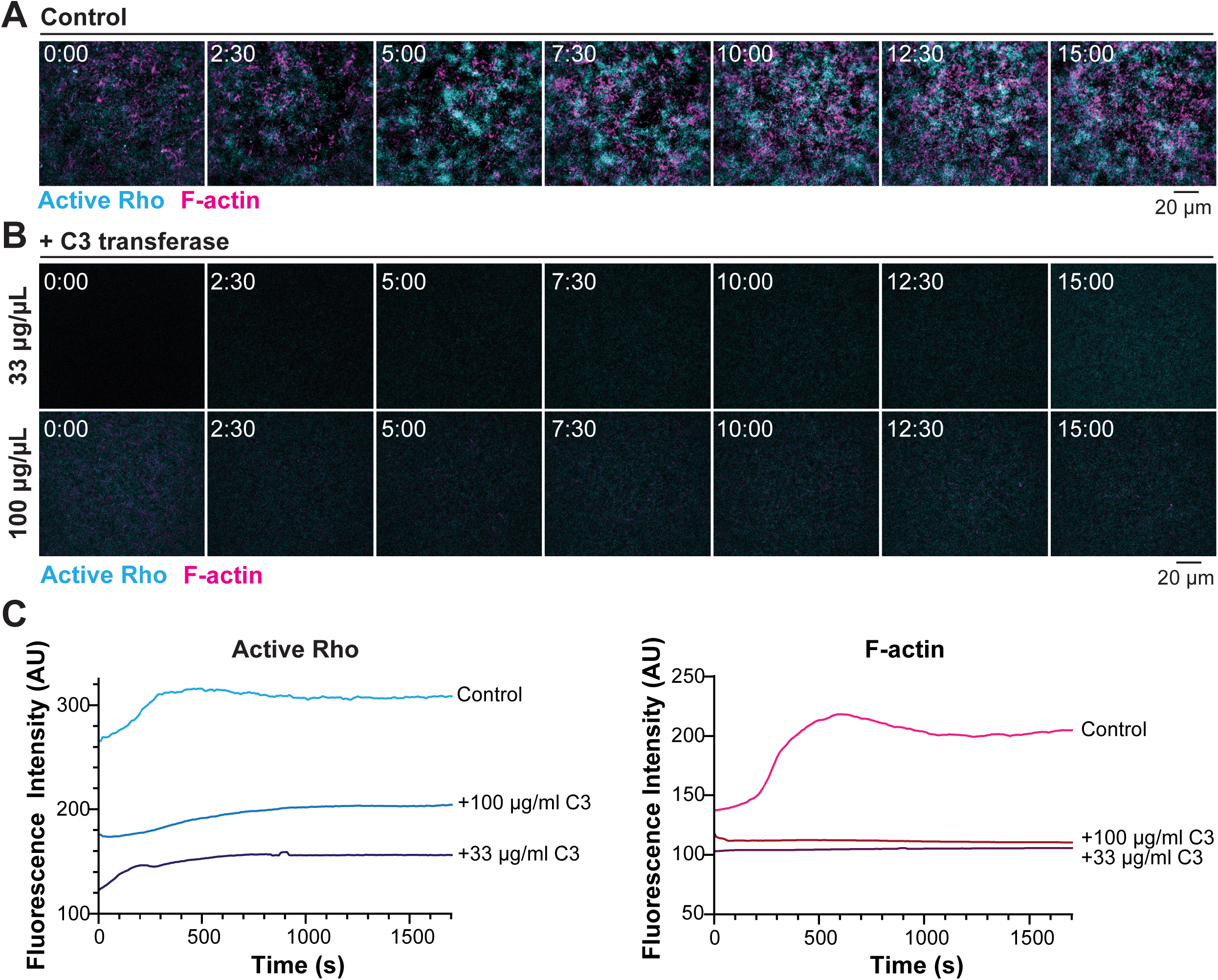
Oscillatory dynamics and waves in the reconstituted cortex require Rho activity. A) Active Rho (cyan) and F-actin (magenta) dynamics in a control cortex. Time is in minutes:seconds. B) Active Rho (cyan) and F-actin (magenta) dynamics following treatment with the Rho inhibitor C3 transferase. Time is in minutes:seconds. C) Quantification of fluorescence intensity of active Rho (cyan) and F-actin (magenta) averaged over the whole field of view in control and C3 transferase-treated extract.

### Reconstituted Rho and F-actin oscillations and waves require polymerized F-actin

We next investigated whether F-actin is required for the emergence of reconstituted cortical dynamics. In starfish embryos, treatment with the actin polymerization inhibitor Latrunculin resulted in a brief increase in Rho wave amplitude as cortical actin was reduced. This was followed by the disappearance of cortical Rho waves when cortical actin was completely lost, leading to the proposal that F-actin is an essential part of the negative feedback in cortical excitability^9^. We therefore allowed oscillations to develop for 30 minutes before adding a vehicle control (DMSO) or 15 μM Latrunculin B (Figure 4A, Supplemental Movie 5). Control, DMSO-treated, extract continued to produce dynamic Rho and F-actin oscillations on the SLB, as observed in kymographs of active Rho and F-actin, whereas in Latrunculin B-treated extract the F-actin and Rho activity oscillations rapidly disappeared (Figure 4B). We quantified the whole field fluorescence intensity of active Rho and F-actin over time and found that in control-treated extract, the amount of active Rho and F-actin associated with the SLB continued to increase after DMSO addition. In contrast, the fluorescence signals of active Rho and F-actin rapidly and completely vanished after Latrunculin B addition (Figure 4C), demonstrating that polymerized F-actin is required for the observed spatiotemporal dynamics. This result differs from the effect of Latrunculin B treatment in starfish embryos, where F-actin disassembly leads to a transient increase in Rho activity before cortical waves are extinguished^9^. However, in both systems, F-actin disassembly leads to the eventual loss of cortical patterning, suggesting that F-actin is an essential regulator of cortical Rho dynamics. These results, taken together with our finding that Rho activity is required for reconstituted dynamics, demonstrates that the molecular mechanisms underlying reconstituted cortical oscillations and waves recapitulate mechanisms underlying cortical waves in cells^9^.

**Figure 4:**
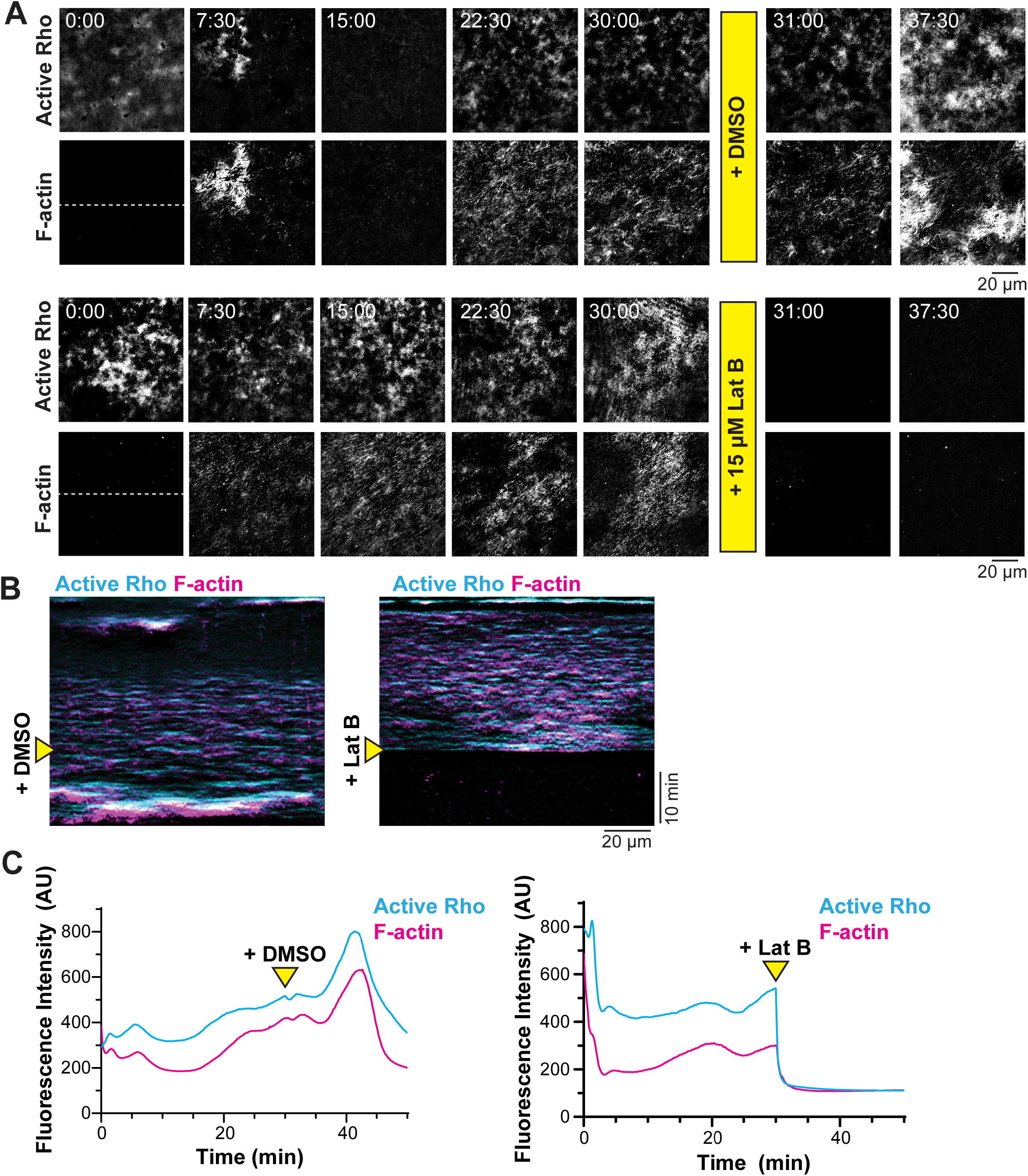
Oscillations in the reconstituted cortex require F-actin polymerization. A) Active Rho and F-actin dynamics before and after addition of DMSO (vehicle control) or 15 μM Latrunculin B. Time is in minutes:seconds. Yellow bars represent the time of DMSO/Latrunculin B addition (30 minutes). Dashed lines represent regions used to generate kymographs. B) Kymographs of active Rho (cyan) and F-actin (magenta) generated from a 10 pixel wide line, as indicated in panel (A). Yellow arrowheads represent the time of DMSO or 15 μM Latrunculin B addition. C) Whole field fluorescence intensity of Active Rho (cyan) and F-actin (magenta) over time in extract treated with DMSO or 15 μM Latrunculin B. Yellow arrowheads indicate time of DMSO or 15 μM Latrunculin B addition.

In addition to Latrunculin B addition, we also sought to suppress cortical F-actin bundles with other actin-destabilizing drugs (Cytochalasin D and Swinholide A) or pharmacological inhibitors targeted against specific actin regulatory proteins including Formins (SMIFH2), Arp2/3 (CK-666), and Rho kinase (H-1152) or recombinant proteins that depolymerize F-actin (cofilin)^19^. In these experiments, we observed an “all or nothing” response, where low concentrations of drugs or cofilin did not prevent F-actin bundles, and high concentrations completely eliminated F-actin association with the SLB (data not shown), similar to the Latrunculin B treatment. This threshold behavior suggests highly nonlinear regulation of cortical F-actin polymerization that provides the right amount of negative feedback needed to enable Rho excitability.

### Conclusions

The experiments presented here provide, to the best of our knowledge, the first reconstitution of oscillatory, Rho GTPase-driven actin dynamics in a cell-free system. The fact that such complex behaviors arise following the combination of cell extract with SLBs directly confirms the ability of the cell cortex to self-organize into complex, dynamic patterns, as originally proposed by E.E. Just more than 80 years ago^1^.

This reconstituted system offers some distinct advantages over cell-based approaches for studying cortical dynamics including the ability to modify the lipid bilayer composition, add recombinant proteins, remove proteins through immunodepletion, and easily add peptide or small molecule inhibitors. Additionally, imaging a 2-dimensional cortex is more straightforward than imaging a 3-dimensional cell, where the cortex changes shape to accommodate cell movment and cytokinesis^8,9,11^.

The reconstituted cortical Rho waves and oscillations shown here exhibit many properties that are similar to cortical waves described in developing embryos^9^; however, there are also clear differences. The excitable, traveling active Rho and F-actin waves identified here do not form a periodic pattern, which we speculate might be due to increased bundled actin accumulation on the SLBs compared to the cell cortex. Highly bundled actin may not provide the right balance of negative feedback to support Rho excitability without suppressing wave propagation. Excess F-actin accumulation on the SLBs following the excitable wave could suppress continued excitability by recruiting a GAP to turn off Rho activity^8,9^. These differences in cortical dynamics between cells and the cell-free system call for further investigation. Identifying the factor(s) that underlie these differences will deepen our understanding of the molecular mechanism that supports self-organization of cortical excitability.

A reconstituted system such as the one presented here could be a powerful approach for studying other types of cortical dynamics including cortical waves that have been identified in migrating cells^6,7^, contractile actomyosin pulses in worm and mouse embryos^4,5^, and subcellular oscillations in non-motile cells^20^. An extract-based reconstituted system also serves as a bridge to a fully reconstituted system consisting exclusively of purified components. Recombinant Rho can dynamically and reversibly interact with a SLB^21,22^, suggesting that investigating fully reconstituted excitability using minimal components could be possible. Overall, cell-free reconstitution of cortical Rho dynamics is an important step towards dissecting the molecular mechanisms of cellular morphogenesis and biological pattern formation and is broadly applicable to a variety of essential cellular functions.

## Abbreviations

(F-actin): Filamentous actin
(SLB): supported lipid bilayer
(rGBD): Rho binding domain of Rhotekin
(UtrCH): Utrophin calponin homology domain
(TIRF): Total Internal Reflection Fluorescence
(PI(4,5)P_2_): Phosphatidylinositol 4,5-bisphosphate
(GAP): GTPase activating protein
(GTP): Guanosine triphosphate

## Acknowledgements

We would like to thank the members of the Miller, Bement, Goryachev and Vecchiarelli labs for their feedback on this research and comments on the manuscript. We thank George von Dassow for his contributions to our thinking about cortical excitability and Kevin Sonnemann and Thomas Burke for their assistance in reagent generation. This work was funded by NSF grants 1615338 (A.L.M.), 1614190 (W.M.B.), and BBSRC grants BB/P006507 and BB/P01190X (A.B.G.). C.M.F. is supported by NIH R35GM131753. J.L. is supported by an American Cancer Society Postdoctoral Fellowship.

## Author Contributions

Conceptualization, J.L., and A.L.M., Methodology, J.L., C.M.F., A.G.V., and A.L.M., Software, M.L. and A.B.G., Validation, J.L. and M.L., Formal Analysis, J.L. and M.L., Investigation, J.L., Resources, A.L.M., A.G.V., M.P., Writing – Original Draft, J.L., M.L., K.S., A.B.G., and A.L.M., Writing – Review & Editing, all authors, Visualization, J.L. and M.L., Supervision, W.M.B., A.B.G., and A.L.M, Project Administration, A.B.G., and A.L.M., Funding Acquisition, W.M.B., A.B.G., and A.L.M.

---

**Supplemental Figure 1**:

A) Left: Schematic of cell-free reconstitution of cortical dynamics. Interphase *Xenopus* egg extract was added to a supported lipid bilayer atop a glass coverslip and imaged using total internal reflection fluorescence (TIRF) microscopy. Active Rho (cyan) and Factin (magenta) associated with the bilayer were visualized using GFP-tagged active Rho probe (rGBD) and Alexa Fluor 647-labeled calponin homology domain of utrophin (UtrCH). Right: Image of the custom aluminum coverslip holder with wells attached. B) Schematic of the approach for analyzing images of reconstituted waves. (1) A 30s (6 frame) difference subtraction was used to reveal the dynamic populations of active Rho (cyan) and F-actin (magenta). This is demonstrated below with a side-by-side kymograph of raw data and difference subtracted data from the same experiment. Cortical dynamics are more easily visualized in the difference subtracted data. (2) The field of view was divided into a grid of boxes each 20×20 pixels. (3) The fluorescence intensity of Active Rho and F-actin was averaged over each box in the field of view. These values were normalized for amplitude to generate time series of fluorescence intensity (see Figure 1C), which were used to determine period and temporal cross-correlation.

**Supplemental Figure 2**:

A) Micrographs of raw data for reconstituted excitable waves of Active Rho and F-actin corresponding to difference subtraction micrographs shown in Figure 1C. B) Histogram of temporal shifts between active Rho and F-actin time series. Temporal shift = 26.2 ± 7.5 s (mean ± standard deviation, n = 1241 boxes from a single experiment). C) Plot of the temporal shift (mean ± standard deviation) between Rho and F-actin in excitable waves calculated for 5 independent experiments. Temporal shift was obtained by statistical analysis of the histogram shown in (B). The number of boxes analyzed for each experiment is indicated on the graph. D) Plot of the excitable wave velocity (mean ± standard deviation) for active Rho across 5 independent experiments. The number of boxes analyzed for each experiment is indicated on the graph.

**Supplemental Figure 3**:

A) Micrographs of raw data for reconstituted oscillatory dynamics of active Rho and F-actin corresponding to difference subtraction micrographs shown in Figure 2A. B) Histogram of temporal shifts between active Rho and F-actin time series. Average temporal shift = 25 ± 5 s (mean ± standard deviation, n = 870 boxes from a single experiment). C) Plot of the active Rho period (mean ± standard deviation) of oscillatory dynamics from 8 independent experiments. The number of boxes analyzed for each experiment is indicated on the graph. D) Plot of the temporal shift (mean ± standard deviation) of oscillatory dynamics from 8 independent experiments. Temporal shift was obtained by statistical analysis of the histogram shown in (B). The number of boxes analyzed for each experiment is indicated on the graph. E) Summary table of wave properties in developing *Xenopus* embryo and reconstituted waves in *Xenopus* egg extract. For excitable waves, values represent mean ± standard deviation from 5 experiments. For oscillatory dynamics, values represent mean ± standard deviation from 8 experiments.

**Supplemental Movie 1**: Reconstituted active Rho and F-actin excitable waves. Experiment 1 relates to Figure 1A and experiment 2 relates to Figure 1C. Difference subtracted (30 seconds) live imaging of active Rho (cyan, left panel) and F-actin (magenta, middle panel) on a SLB. Right panel is a merged image. Time is indicated in minutes:seconds. Playback at 5 frames per second.

**Supplemental Movie 2**: Additional examples of active Rho and F-actin excitable waves. Related to Supplemental Figure 2C and 2D. Difference subtracted (30 seconds) live imaging of active Rho (cyan, left panel) and F-actin (magenta, middle panel) on a SLB. Right panel is a merged image. Time is indicated in minutes:seconds. Playback at 5 frame per second.

**Supplemental Movie 3**: Reconstituted active Rho and F-actin coherent oscillations on an artificial cortex. Related to Figure 2A. Difference subtracted (30 seconds) live imaging of active Rho (cyan, left panel) and F-actin (magenta, middle panel) on a SLB. Right panel is a merged image. Time is indicated in minutes:seconds. Playback at 20 frames per second.

**Supplemental Movie 4**: Reconstituted oscillations and waves depend on Rho activity. Related to Figure 3A and 3B. Difference subtracted (30 seconds) live imaging of active Rho (cyan) and F-actin (magenta) on a SLB in the presence or absence of C3 transferase. Left panel represents control conditions (no C3 transferase added), middle panel shows extract treated with 33μg/ml C3 transferase, and right panel shows extract treated 100 μg/ml C3 transferase. Playback at 10 frames per second.

**Supplemental Movie 5**: Reconstituted oscillations and waves depend on F-actin. Related to Figure 4A. Difference subtracted (30 seconds) live imaging of active Rho (left panels) and F-actin (right panels) before or after the addition of DMSO (top panels) or 15μM Latrunculin B (bottom panels). DMSO and Latrunculin B were added 30 minutes after the start of imaging. Playback at 20 frames per second.

## Methods

### RESOURCE AVAILABILITY

#### Lead Contact

Further information and requests for resources and reagents should be directed to and will be fulfilled by the Lead Contact, Ann L. Miller

#### Materials Availability

Plasmids generated in this study can be obtained from co-author William Bement

#### Data and Code Availability

The data that support the findings of this study are available from the corresponding authors upon reasonable request. Custom code used in this study will be made available at: https://github.com/GoryachevAB-group.

### EXPERIMENTAL MODEL AND SUBJECT DETAILS

Adult *Xenopus laevis* wild type female frogs were purchased from Nasco. Female frogs were injected with human chorionic gonadotropin (HCG) to induce them to lay eggs.

Frogs were housed in a recirculating tank system (Tecniplast), which constantly monitors water quality parameters (temperature, pH, and conductivity) to ensure safe and consistent water quality for an optimal environment for frog health. Daily health and maintenance checks were performed by Animal Care Staff, and frogs were fed frog brittle (Nasco) two times per week.

All studies strictly adhered to the compliance standards of the US Department of Health and Human Services Guide for the Care and Use of Laboratory Animals and were approved by the University of Michigan’s Institutional Animal Care and Use Committee. A board-certified Laboratory Animal Veterinarian oversees our animal facility.

### METHOD DETAILS

#### Preparation of supported lipid bilayers

Powder lipid stocks (Avanti Polar Lipids, see Key Resources Table) were resuspended to working concentrations in chloroform (CHCl3, Sigma). Chloroform-suspended lipids were combined in the following molar ratios: 0.6 PC, 0.3 PS, 0.1 PI (stock A) and 0.6 PC, 0.3 PS, 0.05 PI, and 0.05 PI(4,5)P_2_ (stock B). Lipid stocks were dried using N_2_ gas and kept warm on a 42°C heat block^12^. Dried lipid stocks were vacuum dessicated for 1 hour at 45°C using a centrifugal vacuum (CentriVap, ThermoFisher Scientific). Under N_2_ atmosphere, lipid stocks were resuspended to a final concentration of 5 mM using high-salt extract buffer (HS-XB 200 mM KCl, 1 mM MgCl_2_, 0.1 mM CaCl_2_), vortexed, incubated at 37°C for 30 minutes, and vortexed again. Lipid stocks were then sonicated to generate small unilamellar vesicles (SUVs) using a cup horn sonicator system (700 Watt Sonicator system 110V, QSONICA) in polystyrene tubes at 80W for 10 minutes with pulsed sonication (30 seconds on, 10 seconds off) at room temperature with N_2_ atmosphere. SUVs were sterile filtered using a 0.22 μm filter (Millipore) under N_2_ atmosphere into borosilicate glass vials (ThermoFisher Scientific) and stored at −20°C for less than 3 months.

To prepare supported lipid bilayers (SLBs), SUVs were thawed at room temperature under N_2_ atmosphere. Lipid stocks A and B were mixed at a 49:1 ratio to generate a 0.1% PI(4,5)P_2_ solution^12^. This solution was sonicated for 5 minutes as described above at room temperature and under N_2_ atmosphere. The sonicated SUVs were diluted to 0.5 mM using HS-XB, and CaCl_2_ was added to a 4 mM final concentration. SUVs were added to O_2_ plasma-cleaned chambers (approximately 30 μL per chamber). SLBs were incubated at 42°C for 30 min, and washed three times with three volumes of extract buffer (XB: 100 mM KCl, 1 mM MgCl_2_, 10 mM HEPES, pH 7.7), taking care to maintain a layer of liquid above the SLB^12^. Washed wells were kept at room temp for less than 4 hours before adding extract.

#### Preparation of imaging chambers

Coverglasses (24 x 50 mm No. 1.5, VWR) were washed in a solution of 2% Hellmanex III (Sigma) for 2 hours at 70°C with constant stirring. Coverglasses were rinsed thoroughly with deionized water and allowed to dry overnight in a dust-free chamber until assembly of imaging chambers. A cleaned coverglass was attached to a custom aluminum coverglass mount, comprised of a rectangular slide with a central oval-shaped opening and clamps on either end to hold the coverglass in place (University of Michigan, Scientific Instrument Shop). The cap and lower third of 0.2 mL PCR tubes (Sigma) were removed to create wells for imaging extract^12^. The rim of the PCR tube was dotted with UV glue (Norland Optical Adhesive No. 61, ThermoFisher Scientific) and adhered to the coverglass using 365 nM UV light for one minute. Additional glue was used to reinforce the attachment of the wells, and chambers were exposed to 365 nM UV light for 30 minutes. Chambers were stored in a dust-free container at room temperature until use.

Immediately prior to adding SUVs, imaging chambers were O_2_ plasma cleaned (PE25-JW Benchtop Plasma Cleaning System, PLASMA ETCH) for 10 minutes.

#### Preparation of F-actin intact M-phase Xenopus egg extract

Adult, female *Xenopus laevis* were induced to lay eggs using HCG in Ca^2+^-free Marc’s Modified Ringers (Ca^2+^-free MMR: 100 mM NaCl, 2 mM KCl, 1 mM MgCl_2_, 100 μM EGTA, 5 mM HEPES, pH 7.8)^23^. Eggs were de-jellied in 2% cysteine (Sigma) in Ca^2+^-free MMR, pH 7.8 and rinsed in three volumes of cytostatic factor extract buffer (CSF-XB: XB + 1 μM MgCl_2_, 5 mM EGTA). Lysed or activated eggs were removed as needed. Eggs were washed with CSF-XB plus protease inhibitors (LPC: 10 μg/mL leupeptin, pepstatin A, and chymostatin, all from Sigma) and transferred to a thin-walled UltraClear centrifuge tube (Beckman Coulter). Eggs were packed using a clinical centrifuge at 700 x g for 30 seconds and then 1,400 x g for 15 seconds^12^. Remaining buffer was aspirated using a vacuum, and packed eggs were kept on ice. To crush, centrifuge tubes were transferred to a swinging bucking rotor (TH-660, ThermoFisher Scientific) and centrifuged (Sorvall WX+ Ultracentrifuge, ThermoFisher Scientific) at 15,000 x g for 15 min at 4°C. Cytoplasmic extract was harvested by punching through the side of the tube using an 18G needle and 1 mL syringe and deposited in a 1.5 mL tube on ice^12^. Protease inhibitors (LPC) were added to a final concentration of 10 μg/mL along with energy mix to a final concentration of 9 mM (energy mix: creatine phosphate, 1 mM ATP, 1 mM MgCl_2_, all from Sigma)^12^. This extract is arrested in metaphase of meiosis II^15^.

#### Converting to interphase extract and monitoring the cell cycle state

A controlled release of M-phase arrested extract into interphase (I-phase) is desired. A gelation-contraction assay was used to evaluate the cell cycle state of the isolated extract. Several 2 μL droplets of extract were pipetted under heavy mineral oil (ThermoFisher Scientific) in a petri dish, and light scattering was monitored using a stereoscope. Extract was generally considered high quality if droplets from the (M-phase) control aliquot showed contraction to the droplet center after 5 minutes^15^. If the extract does not contract, the arrest was released during the extract preparation, and therefore, the extract was not used. To convert the extract to interphase (I-phase), CaCl_2_ was added to a single 200 μL aliquot of extract to a final concentration of 0.4 mM, with vigorous mixing^12,15^. Extract was incubated for 5 minutes at room temperature before assessing cell cycle state. Using the gelation-contraction assay, the extract was deemed to be in I-phase if the light scattering particles did not contract toward the center of the droplet after 5 minutes^12^.

#### Preparation of reactions for live imaging

After I-phase cell cycle state was confirmed, extract was placed on ice while reactions were prepared. Recombinant protein fluorescent probes, GFP-rGBD and UtrCH-647 were diluted to working concentrations in freshly-cycled I-phase extract. GFP-rGBD was added to the reaction mixture at a final concentration of 750 nM, and UtrCH-647 was added at a final concentration of 100 mM. After washing with XB, SLBs were washed three times with 30 μL freshly cycled I-phase extract, taking care not to expose the SLBs to air. The reaction mixture was added to the SLB, and the chamber was immediately taken to the microscope for live imaging. Working stocks of GFP-rGBD and UtrCH-647 diluted in I-phase extract were aliquoted, frozen in liquid N_2_, and stored at −80°C.

#### Generation of new DNA constructs

pFastBac1/FLAG-UtrCH was engineered by PCR so that the sequence immediately N-terminal to the Utrophin start site is: M*DYKDDDDK*G**C**G_where *FLAG*-tag is in italics and the lone **cysteine** residue is in bold).

#### Expression and purification of recombinant protein

Baculovirus-mediated recombinant protein expression and anti-FLAG affinity purification were carried out as previously described^9^. Alexa Fluor 647 C2 maleimide was conjugated to the purified protein via the novel cysteine residue according to the manufacturer’s instructions (ThermoFisher).

#### Live imaging

TIRF microscopy was performed using a Nikon Ti2-E motorized inverted microscope with perfect focus, motorized TIRF module, and a LUN-F laser light source (488nm (90mW), 561nm (70mW), and 640nm (65mW)) all controlled by NIS Elements software. A TIRF Quad Dichroic cube (C-FL TIRF Ultra Hi S/N 405/488/561/638 Quad Cube, Z Quad HC Cleanup, HC TIRF Quad Dichroic, in metal cube, HC Quad Barrier Filter) was used with a 60X Objective lens (CFI60 Apochromat TIRF 60X Oil Immersion Objective Lens N.A. 1.49, W.D. 0.12 mm, F.O.V 22 mm) and a Photometrics Prime 95B Back-illuminated sCMOS camera.

Generally, live imaging began within three minutes of completing the buffer and extract washes and adding extract plus recombinant proteins to the SLB. Extract was imaged in the appropriate channel at 5 or 10 second intervals for 60-90 minutes. Perfect focus was used for the duration of the imaging session.

#### Drug treatments

Lyophilized C3 transferase (Cytoskeleton) was resuspended in 1 mM DTT (final buffer: 500 mM imidazole, 50 mM TrisHCl, pH 7.5, 10 mM MgCl_2_, 200 mM NaCl, 5% sucrose, 1% dextran). C3 transferase was aliquoted, frozen in liquid N_2_, and stored at −80°C until use. C3 transferase was added to the extract at a final concentration of 100 μg/mL or 33 μg/mL immediately before extract was added to the SLBs.

Latrunculin B powder (Tocris Bioscience) was resuspended in DMSO (Sigma) to a working concentration of 1 mM and stored at −20°C. For real-time addition of Latrunculin B, an equivalent volume of I-phase extract containing 2X Latrunculin B (final concentration of 15 μM) or DMSO was added to the imaging chamber at the time indicated. Extract containing 2X Latrunculin B was kept at room temperature before adding to the imaging chamber.

### QUANTIFICATION AND STATISTICAL ANALYSIS

#### Figure preparation

Images were processed in Fiji. TIRF mages were cropped to the central portion (approximately one-third of the whole field of view, 674×586 pixels for oscillatory and excitable wave movies). Channels were independently adjusted to highlight relevant features, and LUTs were applied as described in the figure legends. Difference subtraction was performed by duplicating time series stacks with a 30 s time difference (e.g. separated by 6 frames at 5 second interval) and using the “Image Subtraction” tool in Fiji. Kymographs were generated using the Multi Kymograph analysis tool in in Fiji. A 10 pixel wide line was drawn horizontally across the middle of the cropped field of view of a difference subtracted movie. Kymographs were generated for individual channels and later merged. Fiji’s ROI manager was used to ensure fidelity in the placement of the line used to generate kymographs. Kymographs were enlarged 2-fold or 4-fold along the Y axis in Fiji with bilinear interpolation to better highlight separation between active Rho and F-actin signal during oscillations and waves.

#### Analysis of oscillatory dynamics

To avoid optical blur, centrally-located, well-focused subareas of the total field of view (typically 600×600 pixels) were selected for the detailed analysis. Then, the difference subtraction transformation with a delay of n = 6 frames was applied to each image frame as described above (Supplemental Figure 1B, subpanel 1). Negative values were adjusted to 0. Using analysis of simulated wave patterns, we ensured that such transformation does not change either signal periods or the shift between the signals. Then, the field of view was divided into square boxes of 20×20 pixels, and active Rho and F-actin signals were spatially averaged within the boxes (Supplemental Figure 1B, subpanel 2). The size of the box was optimized to provide effective reduction of image noise, yet be sufficiently smaller than the characteristic size of the analyzed patterns. This procedure typically resulted in a matrix of 30×30 boxes per field of view. Next, moving normalization was applied to both active Rho and F-actin signals to remove trends. Every frame was normalized by the maximal and minimal values inside a moving window of 40 frames. Following this initial data preparation, Morlet power spectra (using MATLAB function *cwtft*(*‘Wavelet’,‘mor’*)), autocorrelation, and cross-correlation of active Rho and F-actin signals were computed for each box as described elsewhere^9^. Histograms and cumulative statistics were computed over all boxes. Average periods and their standard deviations were computed from the maxima of Morlet spectra as follows. Morlet spectrum maxima were computed with MATLAB function *findpeaks* for each spatial averaging box and movie frame and then averaged over all time points and boxes.

#### Analysis of solitary excitable waves

First, the immobile fraction of signals was removed by the Fourier transform filter as follows. Temporal Fourier transform was computed (using MATLAB function *fft*) for every pixel of the movie. Then, pixels corresponding to low frequencies were filtered out, and the inverse fast Fourier transform (using MATLAB function *ifft*) was applied. Following this procedure, the field of view was divided into boxes as described above. The shift between the two signals was computed by averaging the cross-correlation function over all boxes. Velocities of waves were determined using kymographs computed on median vertical, median horizontal and two diagonal lines of the field of view. Each line of the kymograph was subdivided into 20 pixel segments and the amplitude of the signal averaged over each segment. For each segment, the maximum value along the time dimension of kymograph was found using MATLAB function *findpeaks*, and the velocity of the wave was computed from the dynamics of thus identified maxima.

#### Whole field intensity measurements

Whole field intensity was measured using difference subtracted images cropped to the central portion of the field of view (734×644 pixels for C3 treatment movies and 636×636 pixels for Latrunculin B movies). Mean gray value was measured over time using the “Measure Stack” tool in Fiji. Intensity vs. time was plotted using Microsoft excel or GraphPad Prism.

## References

1. Just, E.E. (1939). The biology of the cell surface (P. Blakiston’s son & co.).

2. Green, R.A., Paluch, E., and Oegema, K. (2012). Cytokinesis in Animal Cells. Annual Review of Cell and Developmental Biology 28, 29–58.

3. Bement, W.M., Benink, H.A., and von Dassow, G. (2005). A microtubuledependent zone of active RhoA during cleavage plane specification. J Cell Biol 170, 91–101.

4. Michaux, J.B., Robin, F.B., McFadden, W.M., and Munro, E.M. (2018). Excitable RhoA dynamics drive pulsed contractions in the early C. elegans embryo. J Cell Biol 217, 4230–4252. 10.1083/jcb.201806161.

5. Maitre, J.L., Niwayama, R., Turlier, H., Nedelec, F., and Hiiragi, T. (2015). Pulsatile cell-autonomous contractility drives compaction in the mouse embryo. Nat Cell Biol 17, 849–855. 10.1038/ncb3185.

6. Weiner, O.D., Marganski, W.A., Wu, L.F., Altschuler, S.J., and Kirschner, M.W. (2007). An actin-based wave generator organizes cell motility. PLoS Biol 5, e221. 10.1371/journal.pbio.0050221.

7. Iglesias, P.A., and Devreotes, P.N. (2012). Biased excitable networks: how cells direct motion in response to gradients. Curr Opin Cell Biol 24, 245–253. 10.1016/j.ceb.2011.11.009.

8. Goryachev, A.B., Leda, M., Miller, A.L., von Dassow, G., and Bement, W.M. (2016). How to make a static cytokinetic furrow out of traveling excitable waves. Small GTPases 7, 65–70. 10.1080/21541248.2016.1168505.

9. Bement, W.M., Leda, M., Moe, A.M., Kita, A.M., Larson, M.E., Golding, A.E., Pfeuti, C., Su, K.C., Miller, A.L., Goryachev, A.B., and von Dassow, G. (2015). Activator-inhibitor coupling between Rho signalling and actin assembly makes the cell cortex an excitable medium. Nat Cell Biol 17, 1471–1483. 10.1038/ncb3251.

10. Michaud, A., Swider, Z.T., Landino, J., Leda, M. Miller, A.L., von Dassow, G., Goryachev, A.B., Bement, W.M. (2021). Cortical excitability and cell division. Curr Biol. In Press.

11. Nguyen, P.A., Groen, A.C., Loose, M., Ishihara, K., Wuhr, M., Field, C.M., and Mitchison, T.J. (2014). Spatial organization of cytokinesis signaling reconstituted in a cell-free system. Science 346, 244–247. 10.1126/science.1256773.

12. Field, C.M., Pelletier, J.F., and Mitchison, T.J. (2017). Xenopus extract approaches to studying microtubule organization and signaling in cytokinesis. Methods Cell Biol 137, 395–435. 10.1016/bs.mcb.2016.04.014.

13. Benink, H., and Bement, W. (2005). Concentric zones of active RhoA and Cdc42 around single cell wounds. Journal of Cell Biology 168, 429–439.

14. Taniguchi, D., Ishihara, S., Oonuki, T., Honda-Kitahara, M., Kaneko, K., and Sawai, S. (2013). Phase geometries of two-dimensional excitable waves govern self-organized morphodynamics of amoeboid cells. Proc Natl Acad Sci U S A 110, 5016–5021. 10.1073/pnas.1218025110.

15. Field, C.M., Nguyen, P.A., Ishihara, K., Groen, A.C., and Mitchison, T.J. (2014). Xenopus egg cytoplasm with intact actin. Methods Enzymol 540, 399–415. 10.1016/B978-0-12-397924-7.00022-4.

16. Canman, J.C., Hoffman, D.B., and Salmon, E.D. (2000). The role of pre- and post-anaphase microtubules in the cytokinesis phase of the cell cycle. Curr Biol 10, 611–614.

17. Huelsenbeck, J., Dreger, S.C., Gerhard, R., Fritz, G., Just, I., and Genth, H. (2007). Upregulation of the immediate early gene product RhoB by exoenzyme C3 from Clostridium limosum and toxin B from Clostridium difficile. Biochemistry 46, 4923–4931. 10.1021/bi602465z.

18. Just, I., Richter, H.P., Prepens, U., von Eichel-Streiber, C., and Aktories, K. (1994). Probing the action of Clostridium difficile toxin B in Xenopus laevis oocytes. J Cell Sci 107 (Pt 6), 1653–1659.

19. Nishida, E., Maekawa, S., and Sakai, H. (1984). Cofilin, a protein in porcine brain that binds to actin filaments and inhibits their interactions with myosin and tropomyosin. Biochemistry 23, 5307–5313. 10.1021/bi00317a032.

20. Wu, M., Wu, X., and De Camilli, P. (2013). Calcium oscillations-coupled conversion of actin travelling waves to standing oscillations. Proc Natl Acad Sci U S A 110, 1339–1344. 10.1073/pnas.1221538110.

21. Golding, A.E., Visco, I., Bieling, P., and Bement, W.M. (2019). Extraction of active RhoGTPases by RhoGDI regulates spatiotemporal patterning of RhoGTPases. Elife 8. 10.7554/eLife.50471.

22. Graessl, M., Koch, J., Calderon, A., Kamps, D., Banerjee, S., Mazel, T., Schulze, N., Jungkurth, J.K., Patwardhan, R., Solouk, D., et al. (2017). An excitable Rho GTPase signaling network generates dynamic subcellular contraction patterns. J Cell Biol 216, 4271–4285. 10.1083/jcb.201706052.

23. Murray, A.W., and Kirschner, M.W. (1989). Cyclin synthesis drives the early embryonic cell cycle. Nature 339, 275–280. 10.1038/339275a0.

