## Supplemental Figures for "Rho and F-actin self-organize within an artificial cell cortex"

### Supplemental Figure 1

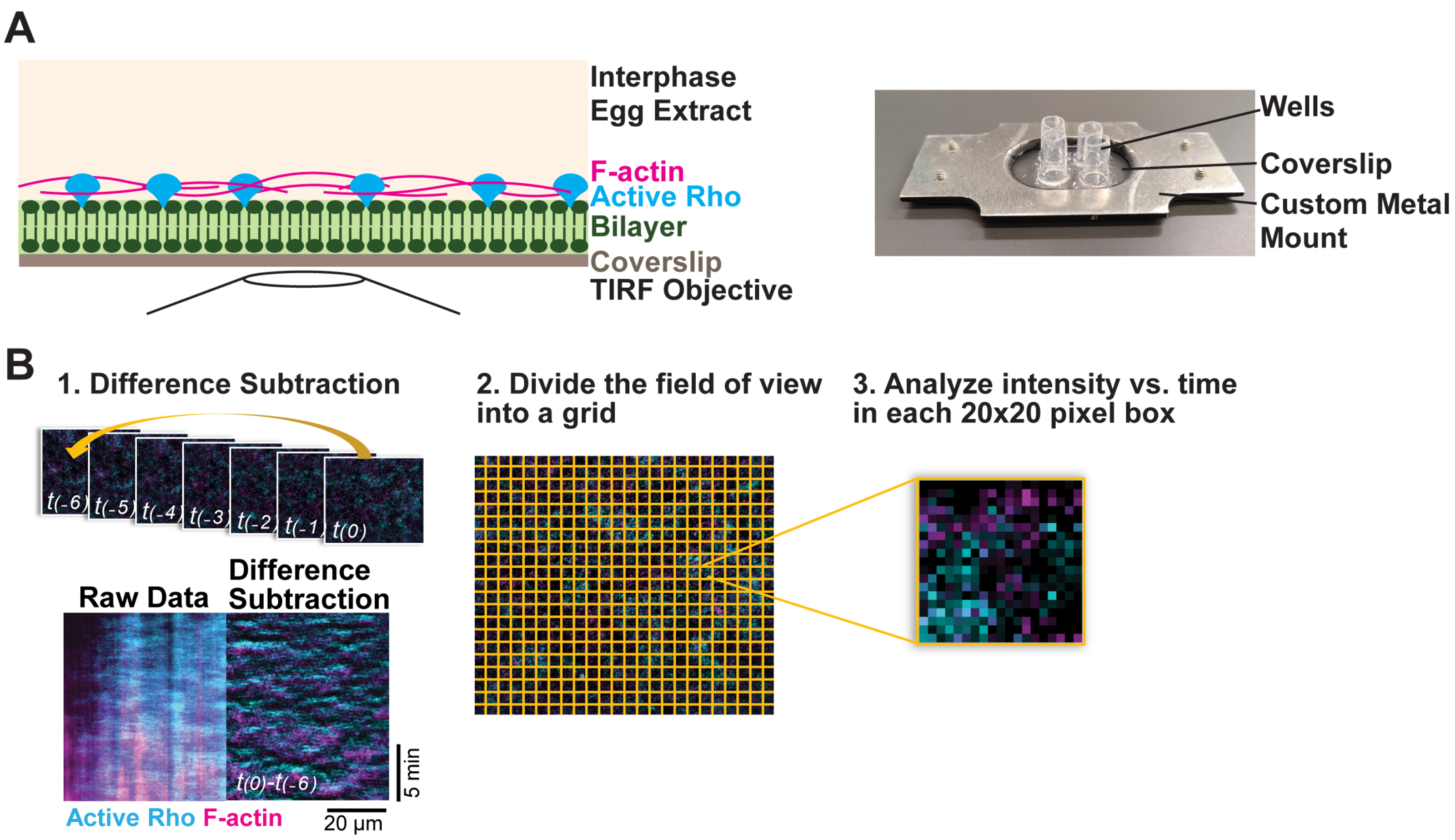

### Supplemental Figure 2

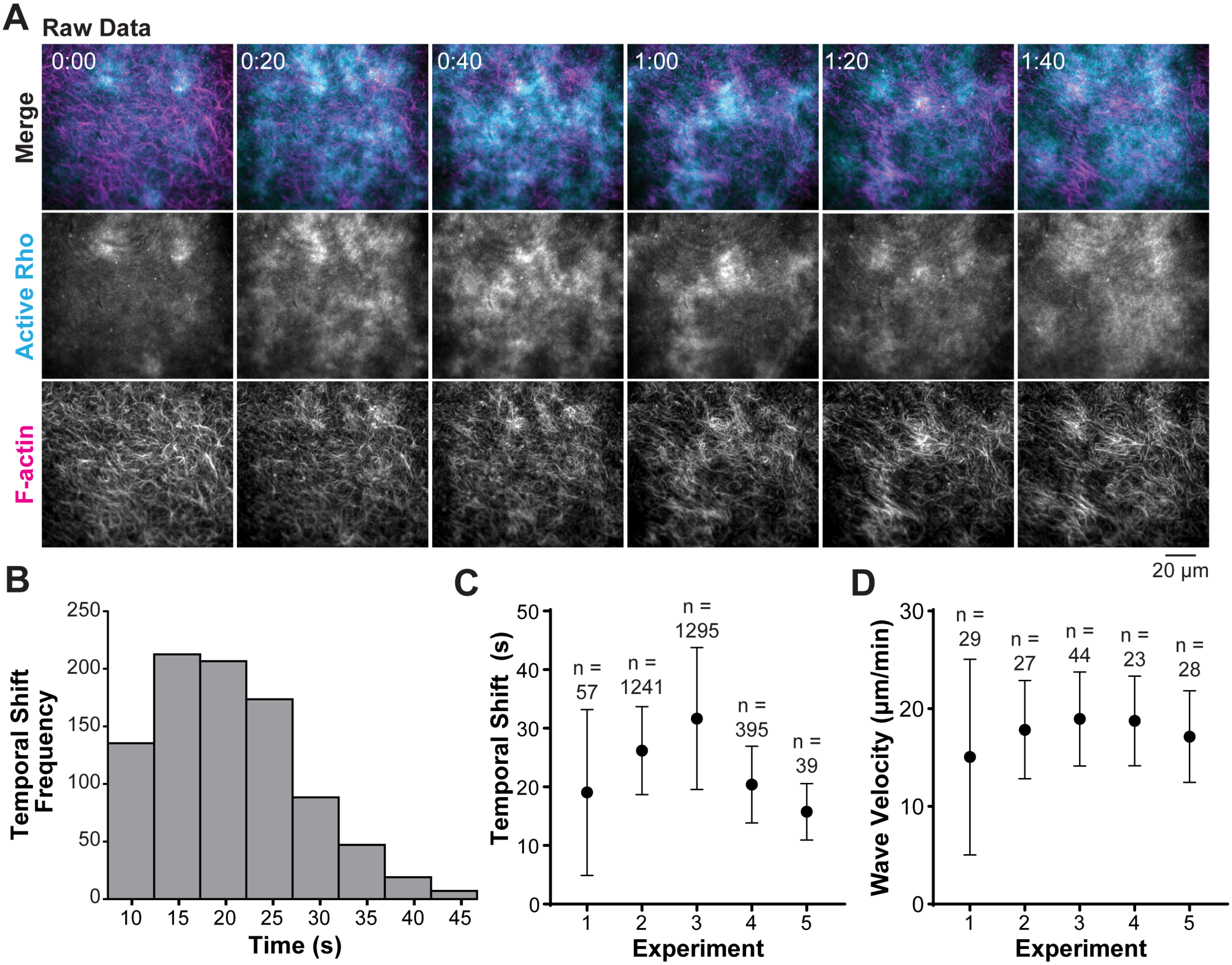

### Supplemental Figure 3

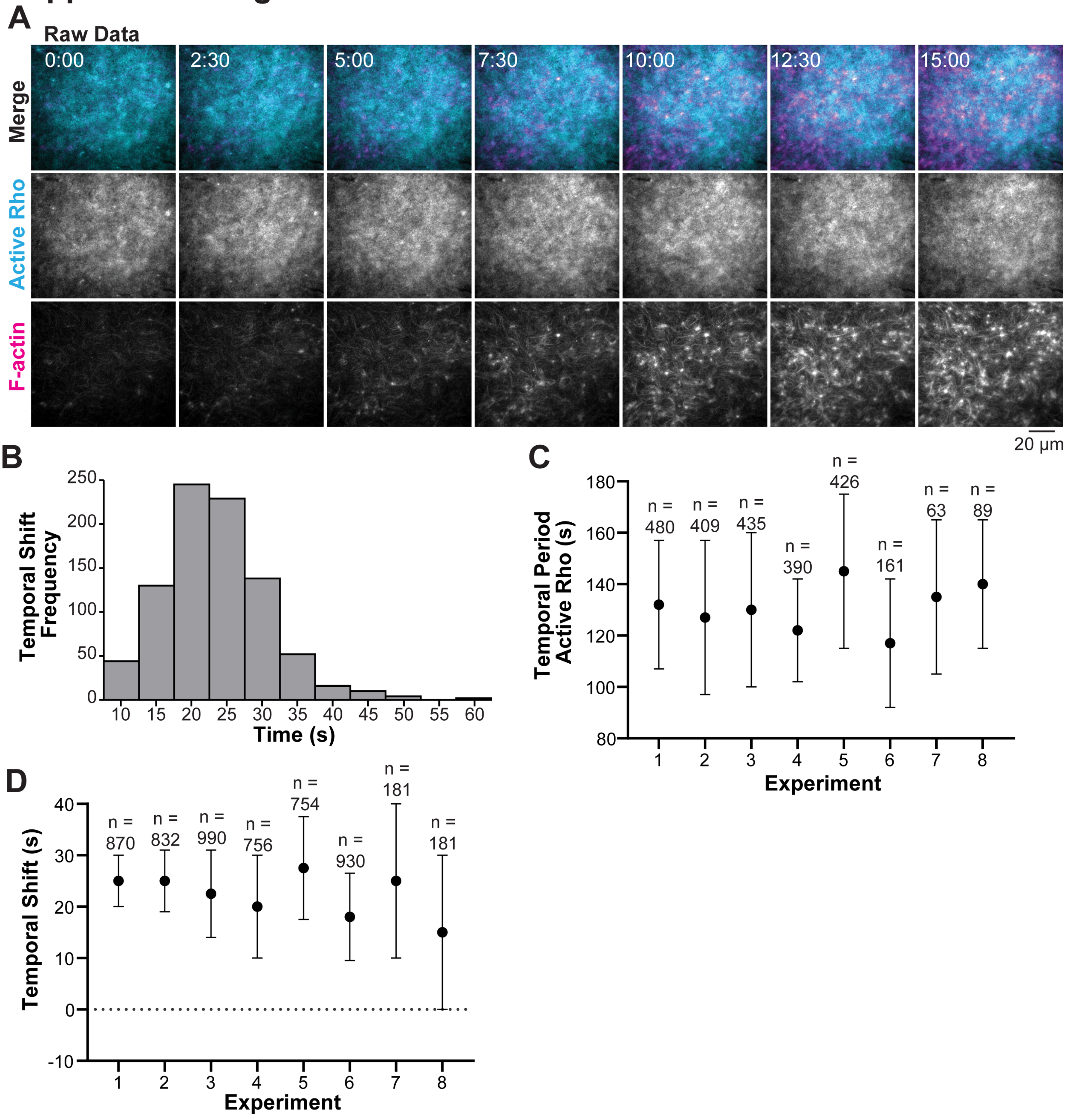


**E**

| | *Xenopus* embryo | *Xenopus* extract: Excitable Waves | *Xenopus* extract: Oscillatory Dynamics |
|---|---|---|---|
| **Time Shift (Cross-correlation)** | 48 seconds | 22.6 ± 6.3 seconds | 22.5 ± 4.1 seconds |
| **Period (Morlet Analysis)** | 80-120 seconds | ——— | 131 ± 8.6 seconds |
| **Velocity** | 13.5 µm/min | 17.6 ± 1.6 µm/min | ——— |
